# Hydrodynamic function of genal prolongations in trinucleimorph trilobites revealed by computational fluid dynamics

**DOI:** 10.1101/2024.01.26.577348

**Authors:** Stephen Pates, Harriet B. Drage

## Abstract

Trilobites, a diverse clade of Palaeozoic arthropods, repeatedly converged on the ‘trinucleimorph’ morphology. Trinucleimorphs possessed vaulted cephala with a broad anterior fringe and prominent posteriorly orientated genal prolongations. Various functional hypotheses have been proposed for the fringe, however the possible function of the genal prolongations has received less attention. Here we use a computational fluid dynamics approach to test whether these prolongations served a hydrodynamic function: generating negative lift to allow trinucleimorphs to remain in place on the seafloor and prevent overturning within fast flowing water. We simulated the performance of cephala with broad, narrow, and absent genal prolongations in a benthic environment with flow speeds ranging from 0.05–0.5 m s^−1^, in two cephalic postures. The first posture had the anterior of the cephalon parallel to the seafloor, while for the second the genal prolongations were parallel to the seafloor. Posture and presence of genal prolongations were found to be important for generating negative lift, with performance improving under faster flow speeds. No significant difference between narrow and broad genal prolongations was detected. This study provides support for genal prolongations serving a hydrodynamic function, similar to the femurs of some insect larvae, however, it does not preclude prolongations also serving additional functions as ‘snowshoes’ or antipredatory deterrents.

## Introduction

Trilobita, a class comprising over 20,000 species, originated in the Cambrian before going extinct at the end of the Permian (Paterson, 2020; Whittington et al., 1997). Representatives of this class are united by the presence of a calcitic dorsal exoskeleton comprising a cephalon, thorax and pygidium, divided laterally into three distinct lobes (Whittington et al., 1997). Trilobites occupied environments across the marine realm, from deep to shallow, some extended into brackish water (Mángano et al., 2021), and filled a wide range of niches, living as, for example, benthic predators, scavengers, filter feeders, pelagic swimmers, and infaunal burrowers (Bergström, 1973; Fortey, 2014, 1985; Fortey and Owens, 1999; Hughes, 2000; Whittington et al., 1997).

The diversity and longevity of trilobites was in part facilitated by the rich array of morphological variation displayed by the group. As trilobite appendages were broadly homonymous in their general structure with limited differentiation (Bergström, 1972; Hughes, 2007, 2003; Losso and Ortega-Hernández, 2022; Pérez-Peris et al., 2021; Walcott, 1918; Whittington, 1975), elaboration and innovation of the dorsal exoskeleton was critical for facilitating the broad ecological disparity of the group (Hughes, 2007).

A number of trilobites converged upon a similar morphology, comprising a vaulted cephalon, broad anterior fringe or brim, and elongate posteriorly orientated genal prolongations, anterior to a short thorax and pygidium (Bergström, 1973; Miller, 1972) (**Figure 1A**). Members of the not closely related trilobite families Dionididae, Harpetidae, Ityophoridae, Brachymetopidae, Raphiophoridae, and Trinucleidae (and rare representatives of other groups) (Chatterton et al., 1994; Fortey and Gutiérrez-Marco, 2022; Fortey and Owens, 1990; Hughes, 2007), are counted among these ‘trinucleimorphs’. Generally considered filter feeders (Adrain et al., 2004; Fortey, 2014; Fortey and Owens, 1999) but see (Pearson, 2017), trinucleimorphs are thought to have been slow moving benthic trilobites that lived on soft substrates (Fortey, 2014; Fortey and Gutiérrez-Marco, 2022; Miller, 1972). The order Trinucleida (Adrain, 2011; Bignon et al., 2020) exemplifies this trinucleimorph form (Hughes et al., 1975), and trinucleids are characteristic members of Ordovician trilobite faunas (see Adrain et al., (2004); distributions in Hughes et al., (1975)). The order Harpida (Beech and Lamsdell, 2021; Ebach and McNamara, 2002) ranged from the late Cambrian (Hughes, 2007) to late Devonian (Lerosey-Aubril and Feist, 2012; McNamara et al., 2009), and were particularly abundant in certain Devonian faunas. While also demonstrating the trinucleimorph form, Harpida tended to have a more flattened cephalic brim in a horseshoe-shape (the ‘harpetid brim’) (**Figure 1B**), this brim sometimes being less pitted than the complex structure of trinucleid fringes (Beech and Lamsdell, 2021; Ebach and McNamara, 2002; Hughes et al., 1975). Harpids also tended to have broader, flatter, and shorter genal prolongations than Trinucleida (Beech and Lamsdell, 2021; Ebach and McNamara, 2002).

**Figure 1.**
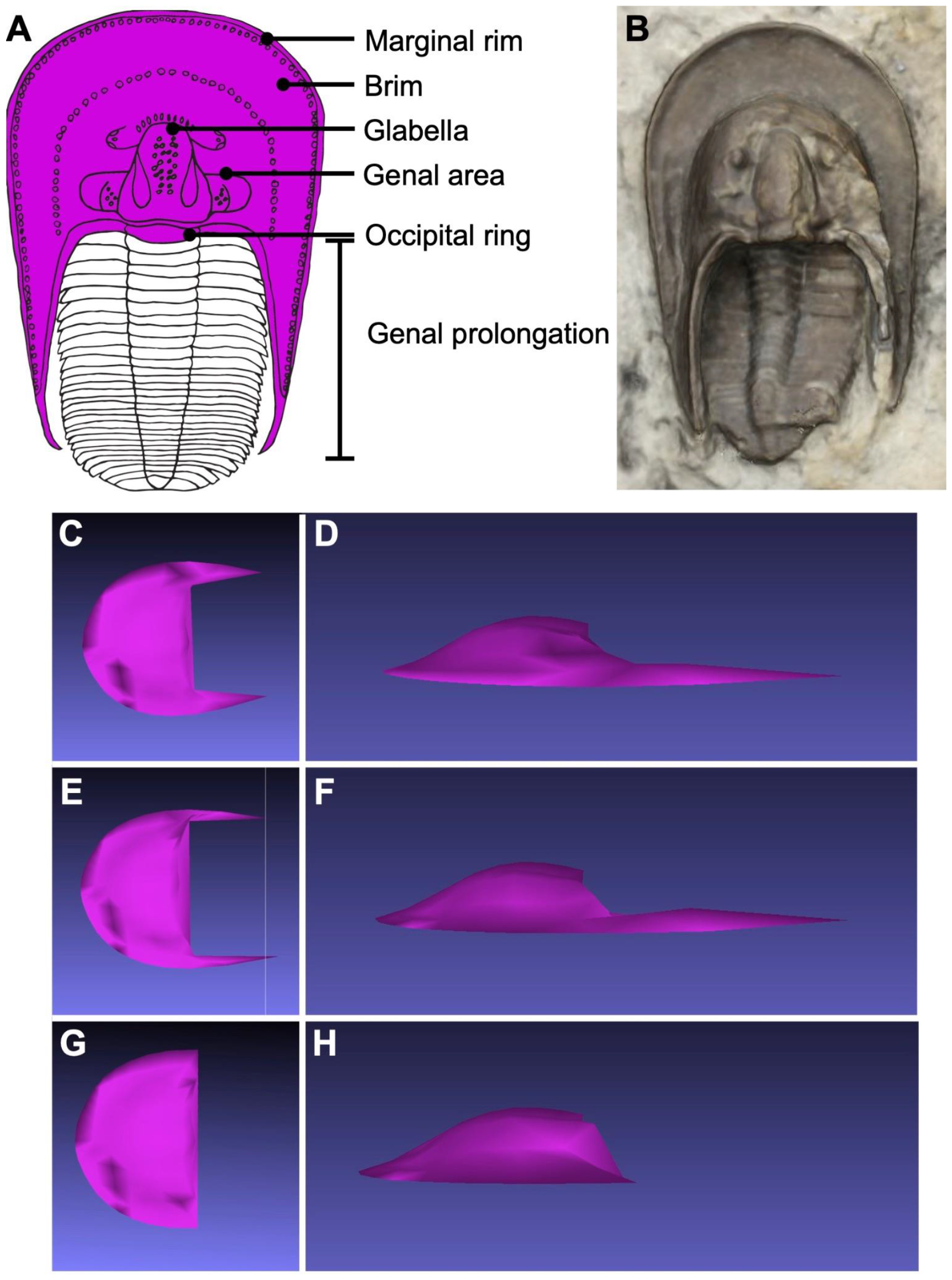
A, representative line drawing of a harpid trilobite, with key morphological features labelled; B, snapshot of 3D model of harpid trilobite, original specimen SMNK-PAL 3891 comes from the Middle Devonian of Auberg bei Gerolstein, Eifel, Germany; C, E, G, dorsal views of the three generalised harpid models used for CFD analyses; D, F, H, lateral views of the three generalised harpid models used for CFD analyses. C and D show the ‘harpid-like’ model with broad genal prolongations; E and F the ‘trinucleid-like’ model with narrow genal prolongations; G and H the model with no genal prolongations.

Harpids were common components of shallow water carbonates, having frequently lived in reef environments throughout the Ordovician to Devonian (Beech and Lamsdell, 2021; Bergström, 1973; Hughes and Thomas, 2014). Members of this group are also found in fine-grained sediments (Miller, 1972), and are known from deeper waters (Beech and Lamsdell, 2021; McNamara and Feist, 2016). In comparison, trinucleids appear to have lived in predominantly deeper waters, having often lost the visual structures (Hughes et al., 1975). Trinucleids are frequently found in black shales and fine sandstones (Miller, 1972; Fortey and Gutiérrez-Marco, 2022), though some might have lived slightly shallower (Waisfeld et al., 2011), or are preserved in distal turbidites (Ghobadi Pour, 2022).

As the morphology has been converged upon multiple times within Trilobita, the adaptive function of the trinucleimorph cephalic shape has drawn much scientific debate. Suggestions generally focus on the prominent fringe, which may have functioned for filter feeding (Bergström, 1972; Fortey and Owens, 1999; Pearson, 2017) and/or burrowing (Bergström, 1972), though Miller (1972) disagrees. The brim of harpids and trinucleids has also been suggested to have functioned like a plough (e.g., Ebach and McNamara, 2002, p. 236), with the latter group suggested to have generated diagnostic trace fossils (Osgood, 1970). The distribution of the pits in the fringe of harpids and trinucleids has also been considered of functional importance (Hughes et al., 1975). Pits may have assisted with sieving, strengthening and/or lightening of the exoskeleton, respiration, or functioned as a sensory organ (e.g., Ebach and McNamara, 2002, p. 236; Miller, 1972), though these hypotheses did not find support when tested experimentally for a single trinucleid species (Pearson, 2017). In comparison, the function of the prominent genal prolongations has received less attention. Genal prolongations in the South American trinucleid *Fantasticolithus isabelae* were suggested to have acted as ‘snowshoes’ that prevented sinking into the substrate (Fortey and Gutiérrez-Marco, 2022), conducive with the ‘mudshoe’ hypothesis proposed for the broad harpid, though not trinucleid, brim (Miller, 1972). Elongate postero-lateral and axial spines in other trilobites have been suggested to perform an antipredatory role (Fortey, 2014; Lynch et al., in press; Pates and Bicknell, 2019; Rustán et al., 2011), particularly when coupled with enrolment behaviour (Chipman and Drage, 2023; Esteve et al., 2011). Indeed enrolment would have orientated the genal prolongations of trinucleids dorsally, in a similar position to some other spines interpreted as antipredatory in many trilobite groups (Fortey and Owens, 1999).

An alternative functional hypothesis for the genal prolongations of trinucleimorphs is rooted in hydrodynamics. Given that trinucleimorphs are considered to have lived in benthic environments, it follows that these individuals required hydrodynamic adaptations that allowed them to ‘stick’ to the seafloor in varied flow regimes, rather than being dislodged by higher velocity ocean currents. In this way the genal prolongations might have functioned similarly to the femurs of mayfly larvae (Ditsche et al., 2023; Weissenberger et al., 1991) or the pectoral fins of bamboo sharks (Wilga and Lauder, 2001). Miller (1972) suggested hypothetical hydrodynamic functions of both the genal prolongations and the brim of harpids. More broadly, investigating the drag and lift forces produced by specific morphological features allows determination of their hydrodynamic impact, if any, as the drag and lift forces an animal is subject to as a result of their size and morphology largely determine the habitat it can survive in (e.g., Vogel, 2008, 1996; Weissenberger et al., 1991).

Here we use a computational fluid dynamics (CFD) approach (e.g., Rahman, 2017) to test whether the elongate genal prolongations in trinucleimorph trilobites played a hydrodynamic role that helped facilitate a benthic lifestyle. We consider the importance of animal posture, and thus orientation of these genal prolongations, on our CFD-simulated outputs.

## Materials and methods

### Experimental design

To determine the impact of trinucleimorph genal prolongations on the hydrodynamic performance of the trilobite cephalon, three sets of simulations were performed (**Figure 1**). The first of these considered a cephalon with broad genal prolongations, the second with no genal prolongations, and the third with narrow genal prolongations more similar to that of trinucleids, rather than harpids. By keeping all other variables constant, including the size, shape, and fringe of the remainder of the cephalon, the impact of the genal prolongations can be isolated. Philosophically this follows the approach of (Shiino et al., 2012), who quantified the performance of the remopleurid trilobite *Hypodicranotus striatulus* with and without the hypostome, to determine the hydrodynamic function of this body part. Importantly, this approach isolates differences in the shape of the genal prolongations from other morphological differences such as the width, orientation and arching of the brim or fringe of trinucleimorphs, or arrangement of pits.

### 3D model creation

A 3D model representing an idealised harpid cephalon was extracted as vertices coordinates from a dataset of trilobite cephala (Pates and Drage, *in prep*). This was then converted to a 3D mesh in Blender. To isolate the impact of the genal prolongations on drag and lift generation, additional models were made by editing this model using Blender to produce identical models except for thinning or removal of the genal prolongations. Blender models were exported as .stl files. These .stl files were imported into ANSYS SpaceClaim. In this software, the function *shrinkwrap* was applied, and then the model was converted to a solid body.

### Computational fluid dynamics virtual flume tank set-up

The SpaceClaim file containing the cephalon model of interest was imported into ANSYS DesignModeler to set up the CFD simulations. Each model was scaled to 25 mm length including the genal prolongations; the model with no genal prolongations was subjected to the same scaling factor but was therefore shorter in total length. Each model was then placed into a virtual flume tank 600 mm long (100 mm in front of the cephalon, 500 mm behind), with a cross-sectional area of 75 × 100 mm (**Fig. 2A**).

**Figure 2.**
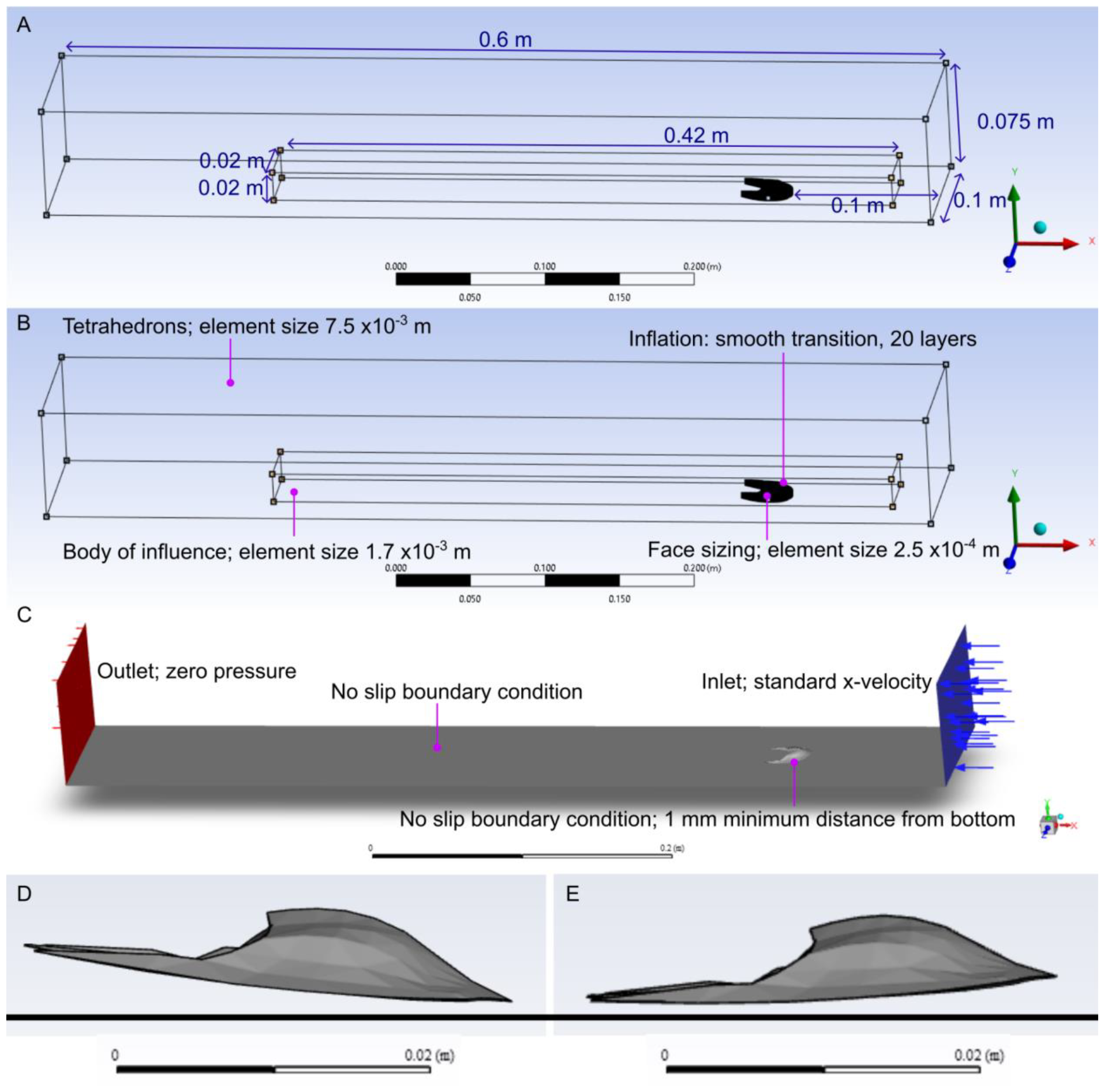
Geometry and meshing set up for ANSYS experiments. A, geometry with dimensions of virtual flume tank, body of influence, and position of model. Model is 0.025 m in length and has a minimum distance of 0.001 m above the bottom of the tank. B, meshing parameters in the simulations for the medium-fine strategy used for the final simulations. Meshing parameters used in mesh refinements given in Supplementary Table 1. C, experimental set-up showing boundary conditions in ANSYS Fluent. D, broad genal prolongations model in anterior horizontal posture. E, broad genal prolongations model in genal prolongations horizontal posture.

The cephala were positioned in two postures for analysis (**Fig. 2D, E**). The first posture had the anterior cephalic margin parallel to the seafloor (comparable to the suggested cephalic orientation of trinucleids of Hughes et al., (1975) and Miller (1972)), and the second had the genal prolongations parallel to the seafloor. For both postures the minimum distance between the ventral surface of the cephalon and the seafloor was set at 1 mm. The first posture simulates a defensive pose, with the genal prolongations pointing dorsally, while the second simulates a pose for horizontal movement (the ‘unstick mechanism’ of Miller (1972)).

### Computational fluid dynamics meshing

The DesignModeler geometry was then passed to ANSYS Meshing. The meshing parameters outlined in **Figure 2B** were used to give an overall mesh of *c*. 2.5 million elements. Meshing refinements of 1.6, 2.1 and 3 million elements were also constructed to test the sensitivity of the results to the number of mesh elements (**Supplementary Table 1**).

### Boundary conditions and model

CFD simulations were run in ANSYS Fluent. Simulations were conducted across a range of inlet velocities (0.05, 0.1, 0.2, 0.3, 0.4 and 0.5 m s^−1^), reflecting flow velocities likely to be encountered by benthic animals (taken from Rahman et al., 2020). An outlet with zero pressure was created at the opposite end. The floor of the virtual flume tank was treated as a no-slip boundary, reflecting a benthic environment; all other boundaries of the tank were treated as frictionless (**Fig. 2C**). The liquid water properties in the ANSYS Fluent Database were used for the fluid (density 998.2 kg m^3^, dynamic viscosity 0.001003 kg m^−1^ s^−1^).

The steady-state pressure-based solver with absolute velocity formulation was chosen for simulations. For the slowest flow speed (0.05 m s^−1^) the laminar model was used. For all other flow speeds (0.1– 0.5 m s^−1^) the realizable k-epsilon model was used with enhanced wall treatment to model the area near the wall.

The Reynolds number (Re) for each simulation can be calculated using equation:

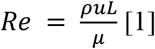

where *ρ* is the fluid density, u is the flow speed, L is the length dimension (here the length of the cephalon is used), and *μ* is the dynamic viscosity of the fluid.

Equation 1 gives Reynolds numbers of 1244, 2488, 4976, 7464, 9952, 12244 for the simulations in this study.

CFD simulations were run for at least 100 iterations, until either all residuals were <1×10^-6^ or 500 iterations had passed. If drag and lift forces on the cephalon had not changed by more than 1/1000 over the final 100 iterations the simulations were considered converged. If after 500 iterations convergence had not been reached simulations were run for another 500 iterations or until convergence was achieved. Drag and lift forces being exerted on the cephalon model were exported at all flow speeds. ANSYS Fluent was used to plot velocity and pressure profiles. Two planes were taken, the first through the axis of the cephalon, the second through the plane of the right genal prolongation.

## Results

For the 2.5 million element mesh, measurements of drag were at most *c*. 2% different from the finer mesh. Differences in lift were at most *c*. 5%, with the exception of the laminar solver simulations used for the lowest flow velocity, where differences in lift values were up to 20%. The laminar solver results were not treated with strong confidence or interpreted further.

The simulated drag and lift forces are presented for both postures in **Figure 3**. With the anterior of the cephalon orientated horizontal to the virtual seafloor, lift forces were negative for all types of genal projection model (**Fig. 3**). With no genal prolongations, the model had lift forces acting on it that were very close to zero (though still negative), and this lift force became gradually more negative with higher flow speeds. In comparison, the other two models (both with genal prolongations) more quickly underwent negative lift forces at increasing flow velocities, always trending at more negative lift than the model with no genal prolongations (**Fig. 3**). The results for the two genal prolongation types (broad, harpid-like and narrow, trinucleid-like) were almost identical for this posture. The models with genal prolongations experienced higher drag forces than the model without prolongations at all flow speeds simulated.

**Figure 3:**
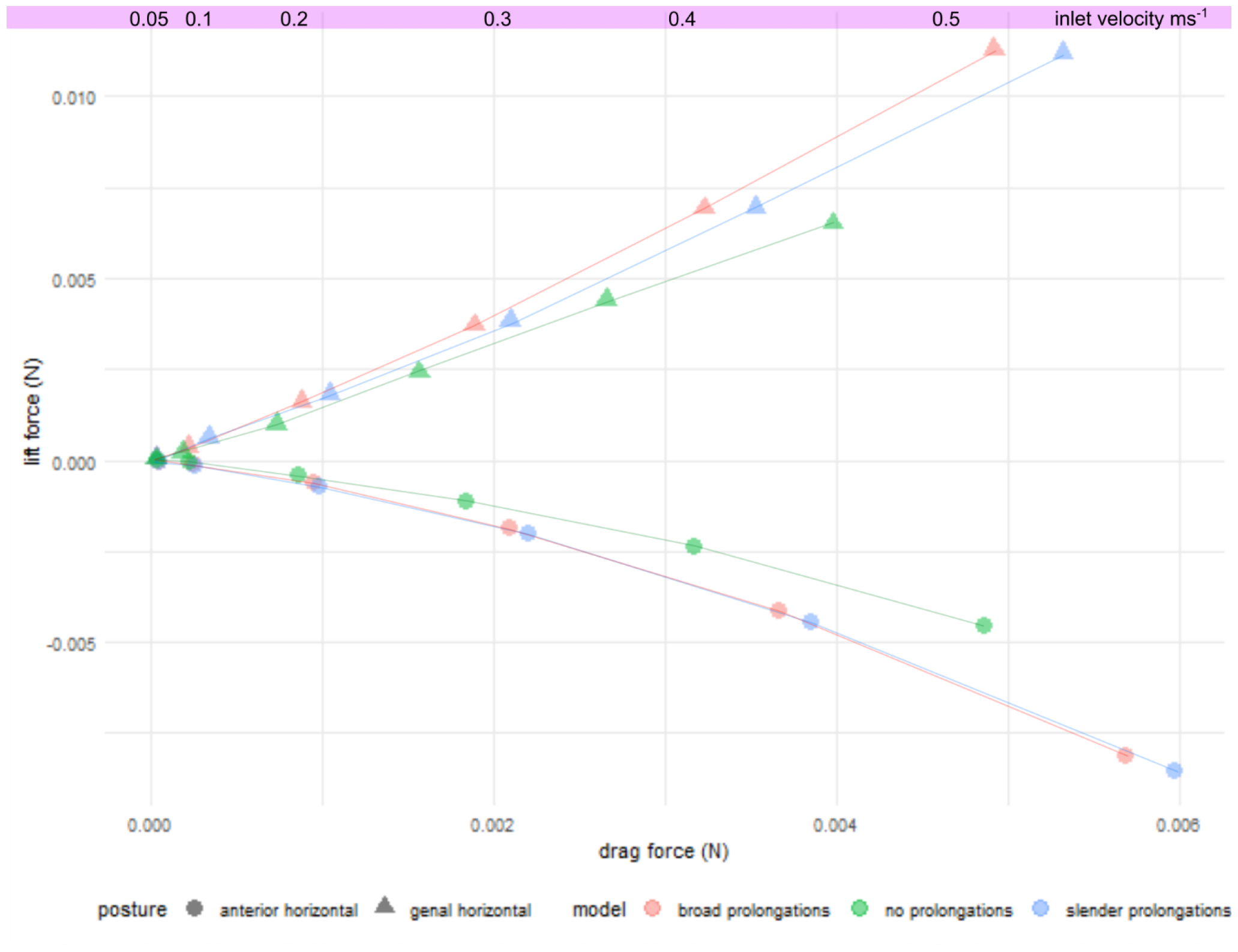
Results of the computational fluid dynamics simulations for all models, both postures, and all flow velocities.

The results show roughly the opposite lift force pattern for the model posture with the genal prolongations positioned parallel to the virtual seafloor (**Fig. 3**). In comparison to the first posture, all lift forces experienced by the models were positive, though lift was lower for the model without genal prolongations. For all models, positive lift increased with an increase in flow speed. The lift force was slightly higher for the harpid-like broad genal prolongations model than the trinucleid-like narrow prolongations model. Again, the drag forces were higher for each flow speed on the two models with genal prolongations compared to the model without (**Fig. 3**).

Static pressure contours were visualised for the models with harpid-like broad genal prolongations (**Fig. 4**), trinucleid-like narrow prolongations (**Fig. 5**) and no prolongations (**Fig. 6**). The pressure projection onto the plane at the midline of the models does not show much variation (**Figs. 4A, 5A, 6A**), with higher pressure at the anterior of the cephalon and low pressure postero-dorsally. A larger pressure difference can be seen between the models with genal prolongations (**Figs. 4B, 5B**) and model without genal prolongations when projected at the plane through the right-hand prolongation (**Fig. 6B**). For the former two models, an area of low pressure extends from midway along the ventral margin of the cephalon and continues beneath the genal prolongation (**Figs 4B, 5B**). For the model without genal prolongations this area of low pressure abruptly terminates at the posterior of the cephalon (**Fig. 6B**). The velocity vectors (**Figs. 7–9**) show similar results for the planes through the midline, with an area of low velocity recirculation immediately posterior to the cephalon. The plane through the right genal prolongation shows no recirculation posterior to the prolongations, and an area of lower velocity beneath the prolongation, with vectors aligned antero-posteriorly (**Figs. 7B, 8B**). Velocity vectors are instead parallel to the ventral surface of the flume tank in the model when there is no genal prolongation (**Fig. 9B**).

**Figure 4.**
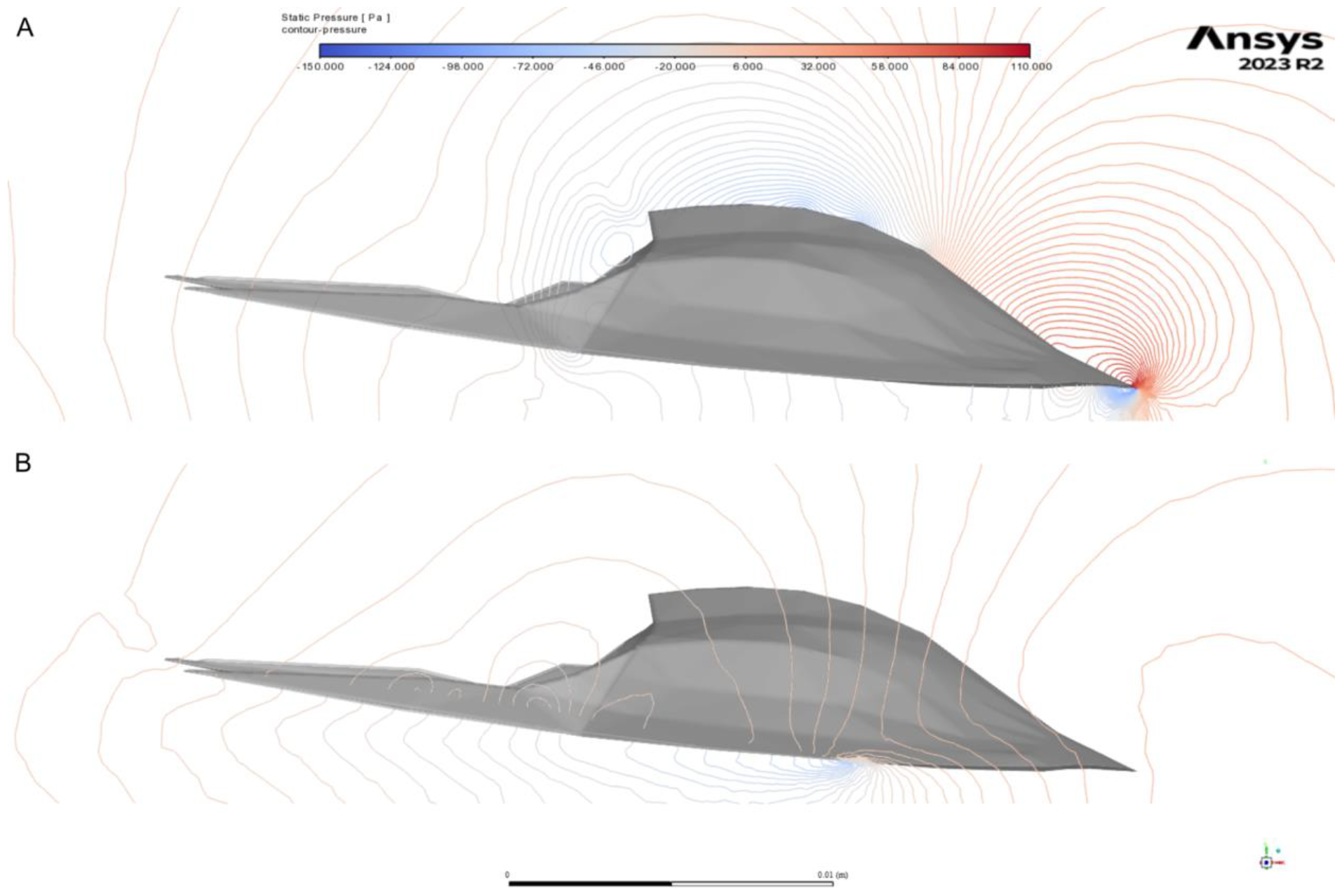
Pressure contours around the model with broad genal prolongations. A, visualised through the midline; B, visualised through the right genal prolongation.

**Figure 5.**
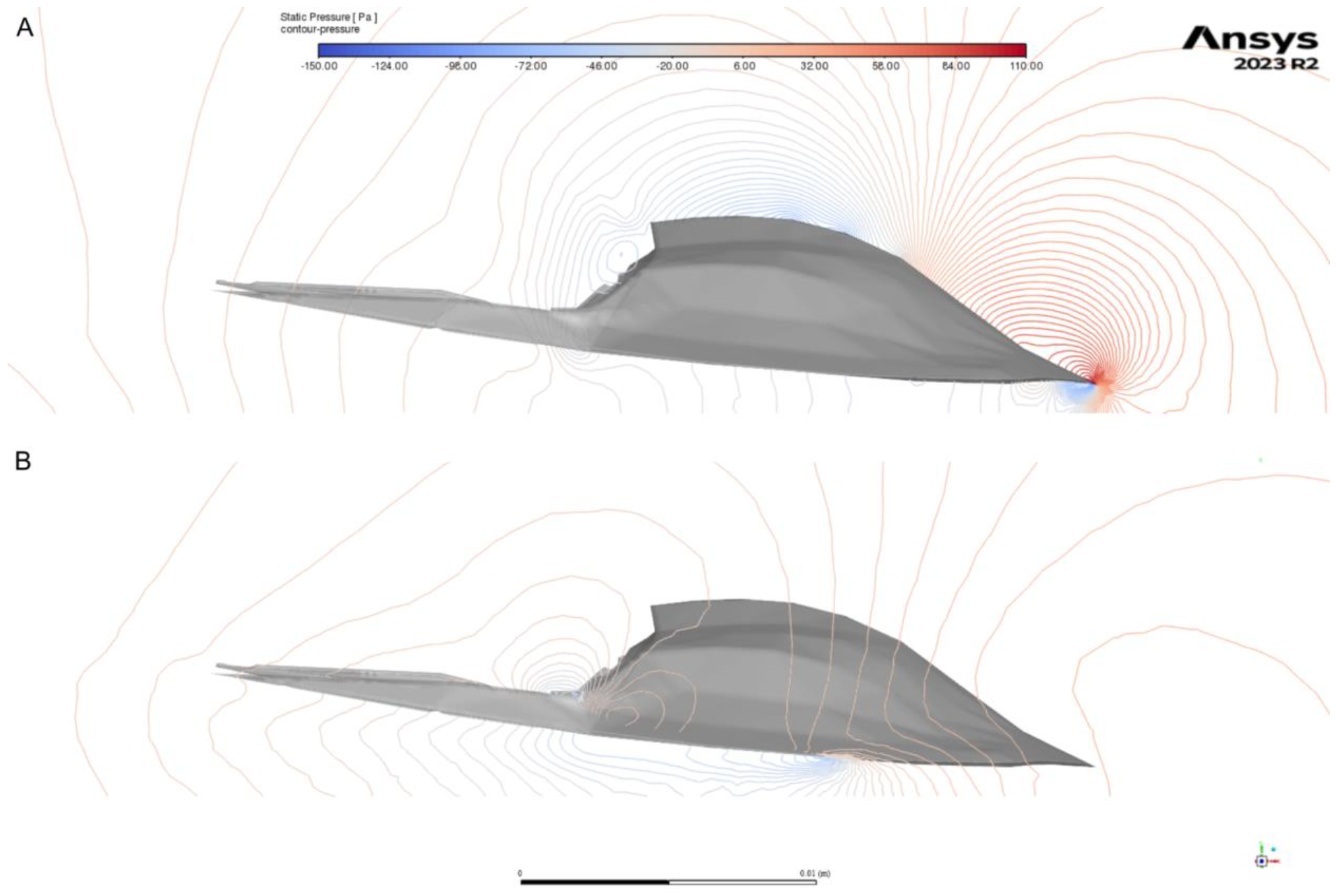
Pressure contours around the model with narrow genal prolongations. A, visualised through the midline; B, visualised through the right genal prolongation.

**Figure 6.**
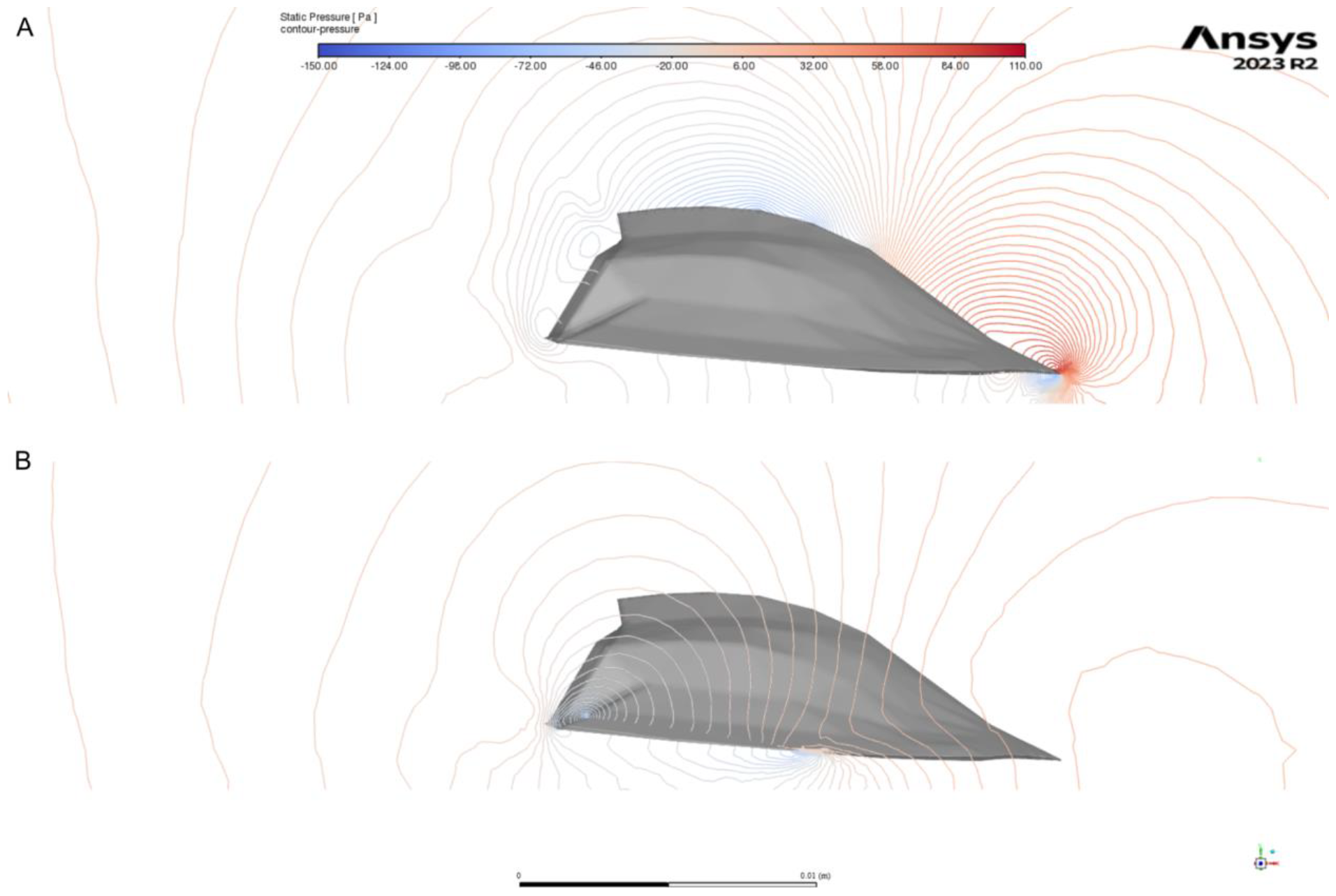
Pressure contours around the model without genal prolongations. A, visualised through the midline; B, visualised through the right genal prolongation.

**Figure 7.**
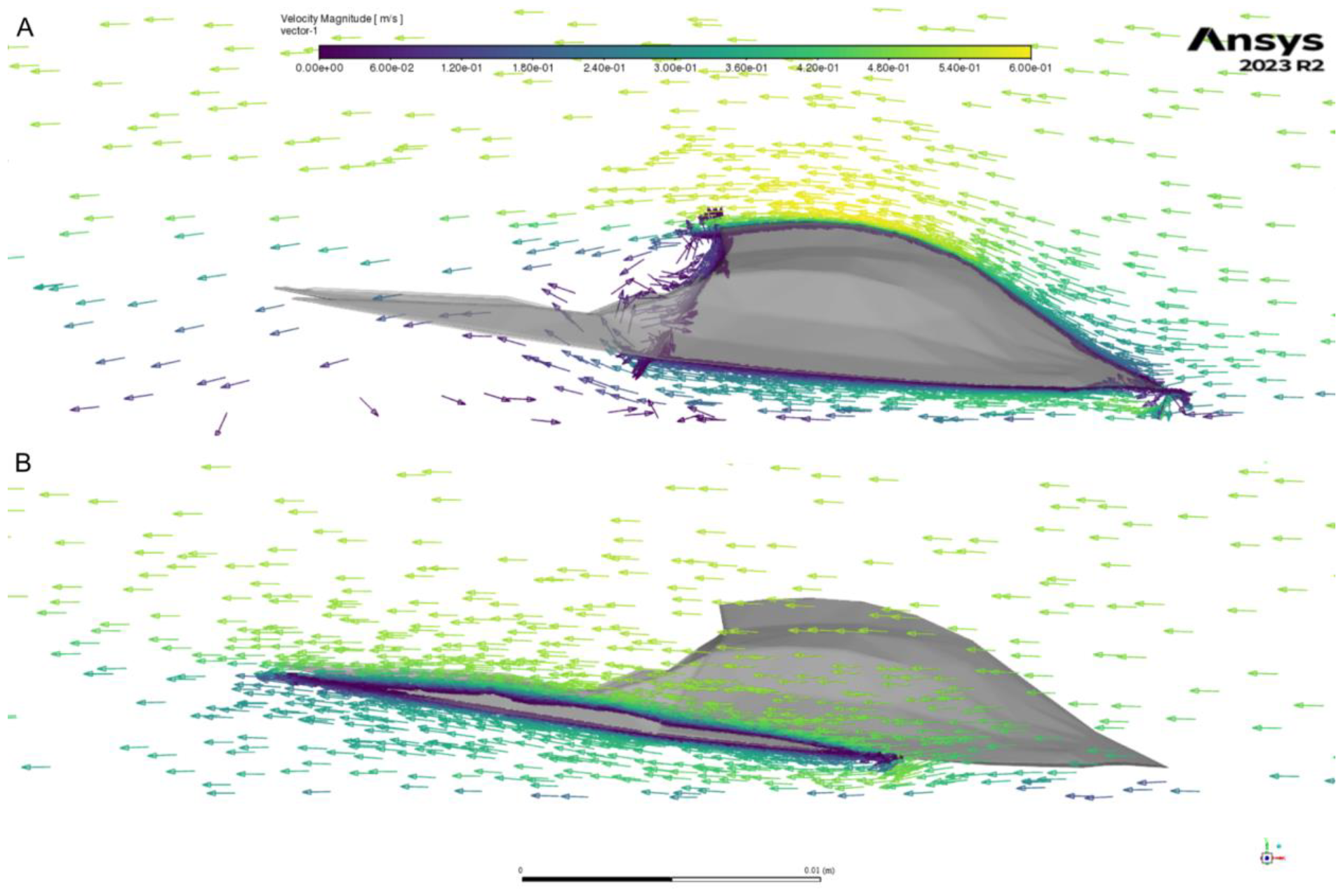
Vectors of velocity around the model with broad genal prolongations. A, visualised through the midline; B, visualised through the right genal prolongation.

## Discussion

Comparison of the hydrodynamic performance of the different models demonstrates that genal prolongations could have served to produce downforce and help prevent benthic trinucleimorph trilobites from being displaced or flipped over during periods of fast water flow (Miller, 1972). This is supported by the more negative lift forces consistently acting on the models with genal prolongations (harpid- and trinucleid-like) compared to no prolongations (**Fig. 3**). This function would have been useful in particular for harpids living in shallow marine onshore environments (e.g., *Scotoharpes* (Beech and Lamsdell, 2021; Fortey and Wilmot, 1991)), which are expected to have been more impacted by storms, have faster and more unpredictable flow velocities, and thus stronger velocity gradients. In such habitats, the danger of being dislodged would be influenced more by lift forces than drag forces, favouring morphologies that generate negative lift (Weissenberger et al., 1991). While all models were able to generate negative lift in the horizontal anterior posture (**Fig. 3**), the presence of elongate genal prolongations generated an additional 100% negative lift force at the highest flow velocity simulated, with a corresponding *c*. 20% increase in drag force (**Fig. 3**).

Comparison of the pressure and velocity projections in the plane through the genal prolongation (**Figs. 4–9**) demonstrates that this downforce is created by the production of an area of low pressure and change in flow direction (orientated slightly upwards, parallel to the genal prolongations) beneath the prolongations. The slender (trinucleid-like) and broad (harpid-like) prolongations produced very similar results (**Figs. 4, 5, 7, 8**), however, the model lacking prolongations did not create such an area of low pressure, nor did it change the direction of the flow, and so this model produced less negative lift values (**Figs. 6, 9**). The production of negative lift comes with the trade-off of increased drag forces, and so such an adaptation would be favoured for slower moving trilobites, as suggested for harpids by Miller (1972), that fed by filtering or sediment sifting. For more active trilobites, such as those pursuing prey, the increased drag costs of negative lift production might have been too high. Interestingly, trinucleids have been suggested to have been more active than harpids based on their fringe morphology (Miller, 1972), but the comparable results for the harpid and trinucleid-like models do not bear this out for the genal prolongation morphology alone. Future studies interrogating the hydrodynamic implications of differences in the morphology of the brim or fringe between harpids and trinucleids (such as degree of vaulting, presence of anterior archways, or even pattern of anterior pits) are required to fully interrogate this hypothesis.

**Figure 8.**
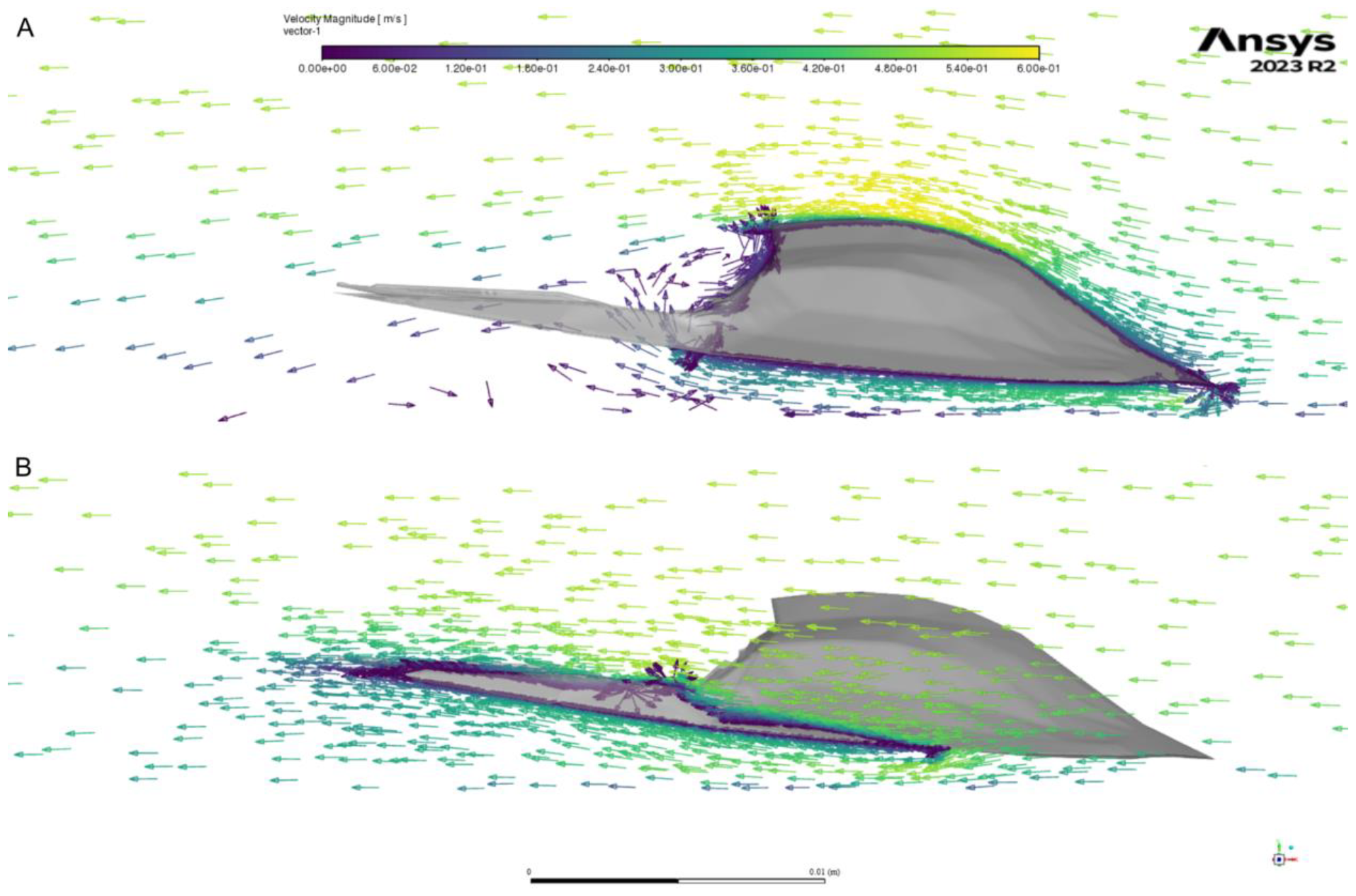
Vectors of velocity around the model with narrow genal prolongations. A, visualised through the midline; B, visualised through the right genal prolongation.

**Figure 9.**
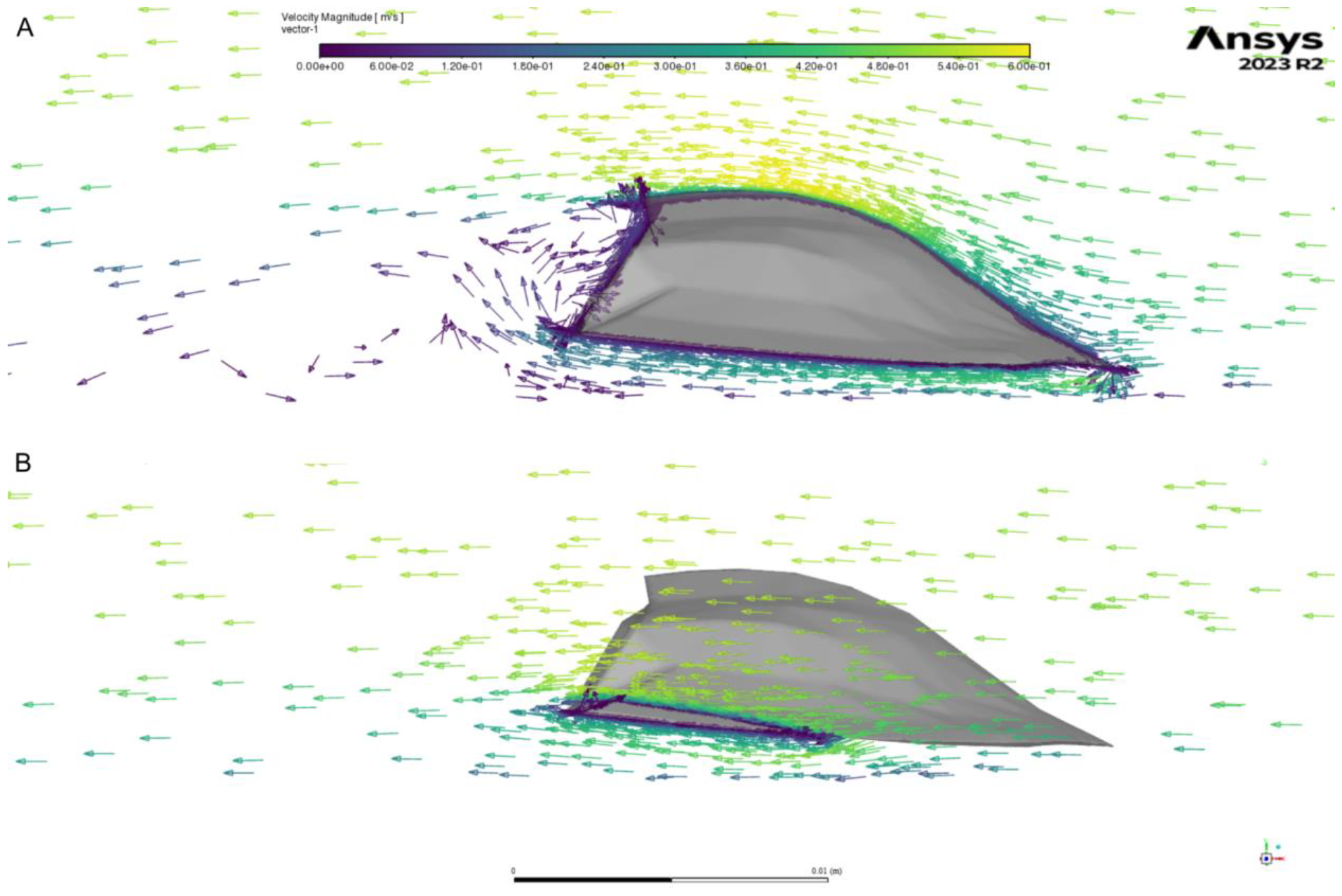
Vectors of velocity around the model without genal prolongations. A, visualised through the midline; B, visualised through the right genal prolongation.

The simulated negative lift force production is comparable to that generated by some extant marine and freshwater arthropods. In simulations of flow around the body of the horseshoe crab *Limulus polyphemus* the orientation of the body and telson facilitated a change in direction of the flow (Davis et al., 2019) comparable to that produced by the genal prolongations in this study. For mayfly larvae, which encounter similar Reynolds numbers to those simulated in this study albeit in a freshwater rather than marine environment, the orientation of the femur appears to serve a similar function to that hypothesised for genal prolongations here; orientating the femurs to direct the flow can generate negative lift and prevent animals being swept away by fast water flow (Ditsche et al., 2023; Weissenberger et al., 1991).

When the genal prolongations were orientated parallel to the seafloor, flow was not redirected and no negative lift was produced. The higher drag force and larger surface area led to greater positive lift forces in the models with broader prolongations (**Fig. 3**). Posture and orientation of the body or body parts in the flow has been shown to be important for generating positive and negative lift, and minimising drag, for other benthic arthropods such as *L. polyphemus* (Davis et al., 2019) and mayfly larvae (Weissenberger et al., 1991). By changing the orientation of the cephalon in the flow, a trilobite would have been able to go from a negative to a positive lift regime, depending on the exact conditions they were experiencing at the time. A small amount of positive lift may have been useful for counteracting the negative buoyancy of the body, while negative lift would have been useful for preventing overturning or dislodgement during storms, and possibly also for facilitating more rapid turns while moving on the seafloor. The danger of being dislodged or overturned is especially pertinent for animals living in environments with large velocity gradients (Weissenberger et al., 1991). For *L. polyphemus* the telson facilitates righting after being inverted (Davis et al., 2019). However, with their generally short thoraces and lack of an elongate spinose process on the pygidium, trinucleimorph trilobites would not have been able to right themselves in a similar way, making the generation of negative lift and prevention of inversion all the more pertinent for trinucleimorphs living in shallower water accompanying larger velocity gradients.

Importantly, the hydrodynamic function of the genal prolongations described above does not preclude the existence of other adaptive functions. For example, the ‘snowshoe’ hypothesis (Fortey and Gutiérrez-Marco, 2022) suggests that, thanks to the genal prolongations, trinucleimorphs with broader genal prolongations were less prone to sinking into the sediment due to their larger surface area. These genal prolongations may have also served as predatory deterrents, like for other trilobite spines (Fortey and Owens, 1999; Pates and Bicknell, 2019). In support of an antipredatory function, harpids with repaired genal prolongations have been reported in the literature (Sinclair, 1947, figs. 1, 2), alongside trinucleimorphs with repaired injuries to the fringe, some of which were inflicted by predators when the animal was enrolled to defend itself (Snajdr, 1979). However, for other repaired injuries to the trinucleimorph fringe, the evidence is more equivocal and self-injury during moulting has been a preferred explanation (Owen, 1985).

## Conclusion

Computational fluid dynamics (CFD) simulations across a range of flow velocities suggest that genal prolongations of trinucleimorphs facilitated the generation of negative lift (downforce), when the cephalon was orientated with the anterior parallel to the seafloor. This function was served by both slender (trinucleid-like) and broad (harpid-like) genal prolongations, and was more effective under faster flow velocities. This result suggests that the genal prolongations were useful for preventing lifting off the bottom or overturning of benthic trinucleimorph trilobites. When the cephalon was orientated with the genal prolongations parallel to the seafloor, the genal prolongations produced more positive lift at faster flow velocities. This may have been useful for locomotion, counteracting negative buoyancy of the body. The genal prolongations, while having a hydrodynamic function, may also have been adaptive in other ways, including preventing sinking into fine sediment (snowshoe hypothesis) and as a predatory deterrent or defence. CFD methods such as that presented here, testing hypothetical models that differ in only one key morphological structure to analyse the structure’s hydrodynamic function, can be applied to a suite of morphological features as putative adaptations to a marine life. This approach builds on and complements previous studies using CFD to address functional and ecological hypotheses relating to trilobites for over a decade (Esteve et al., 2021; Esteve and López-Pachón, 2023; Hou et al., 2023; Shiino et al., 2014, 2012; Song et al., 2021), and previous experimental fluid dynamics approaches using physical models in flume tanks (Fortey, 1985; Lask, 1993).

## Acknowledgements

We thank Lukáš Laibl (Czech Academy of Sciences) and Sarah R Losso (Harvard University) for discussions on harpid morphology and function. Julien Kimmig (Staatliches Museum für Naturkunde Karlsruhe) kindly facilitated study of material in the SMNK-PAL collection, and provided equipment for photogrammetry setup used to create model in Figure 1B.

## Data availability

Spreadsheets with results from CFD simulations and R code used to make Figure 3 are available through the Open Science Framework (doi.org/10.17605/OSF.IO/2MAFU). Unfortunately, the size of the ANSYS projects (as archives or in raw form) exceeds the maximum file size that can be uploaded to the OSF (5Mb). Please contact the authors for a copy of these projects (will be sent as an ANSYS 2023 R2 archive .wbpz file).

## Author contributions

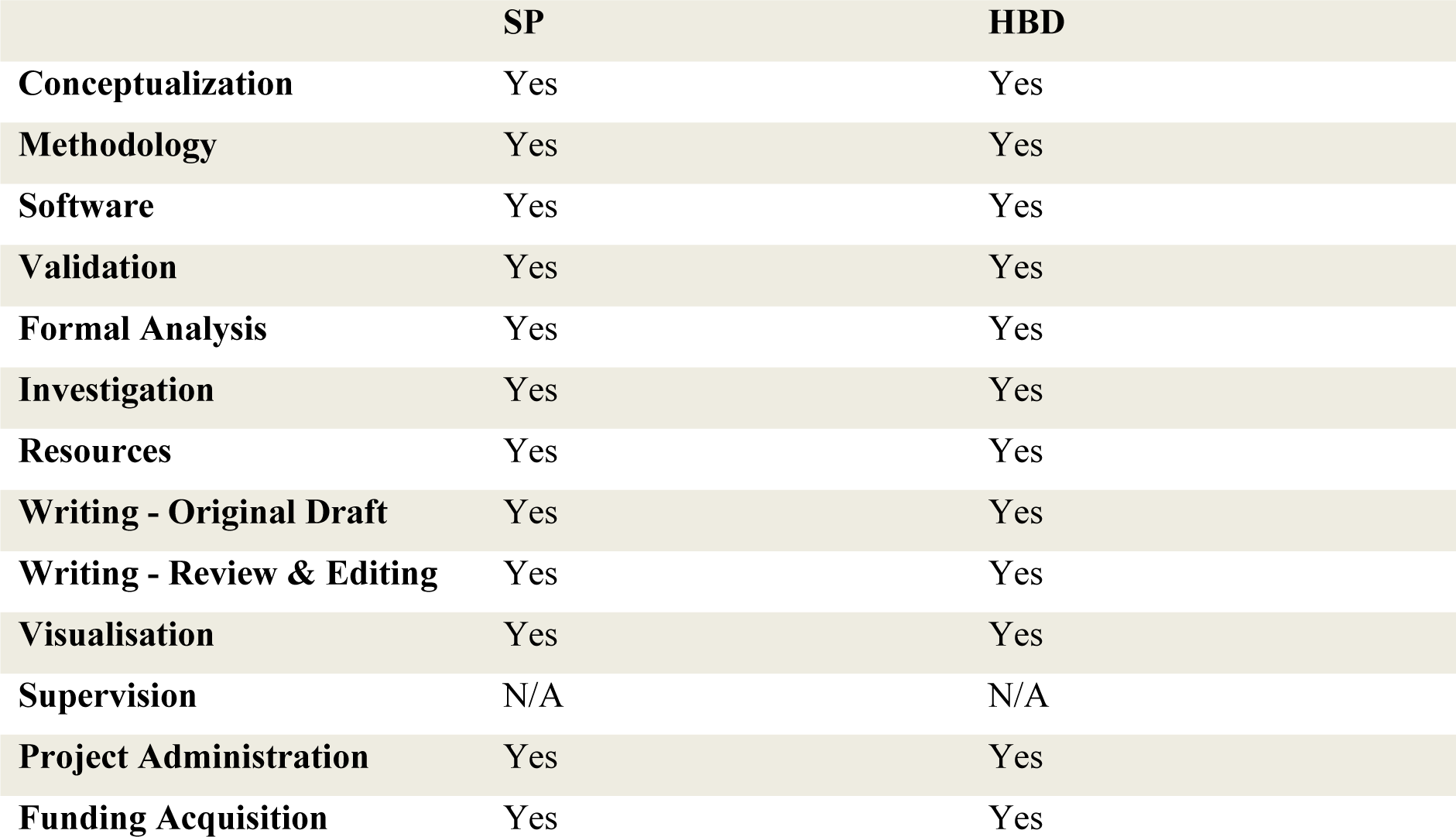

## Supplementary figure and table

**Supplementary Figure 1.**
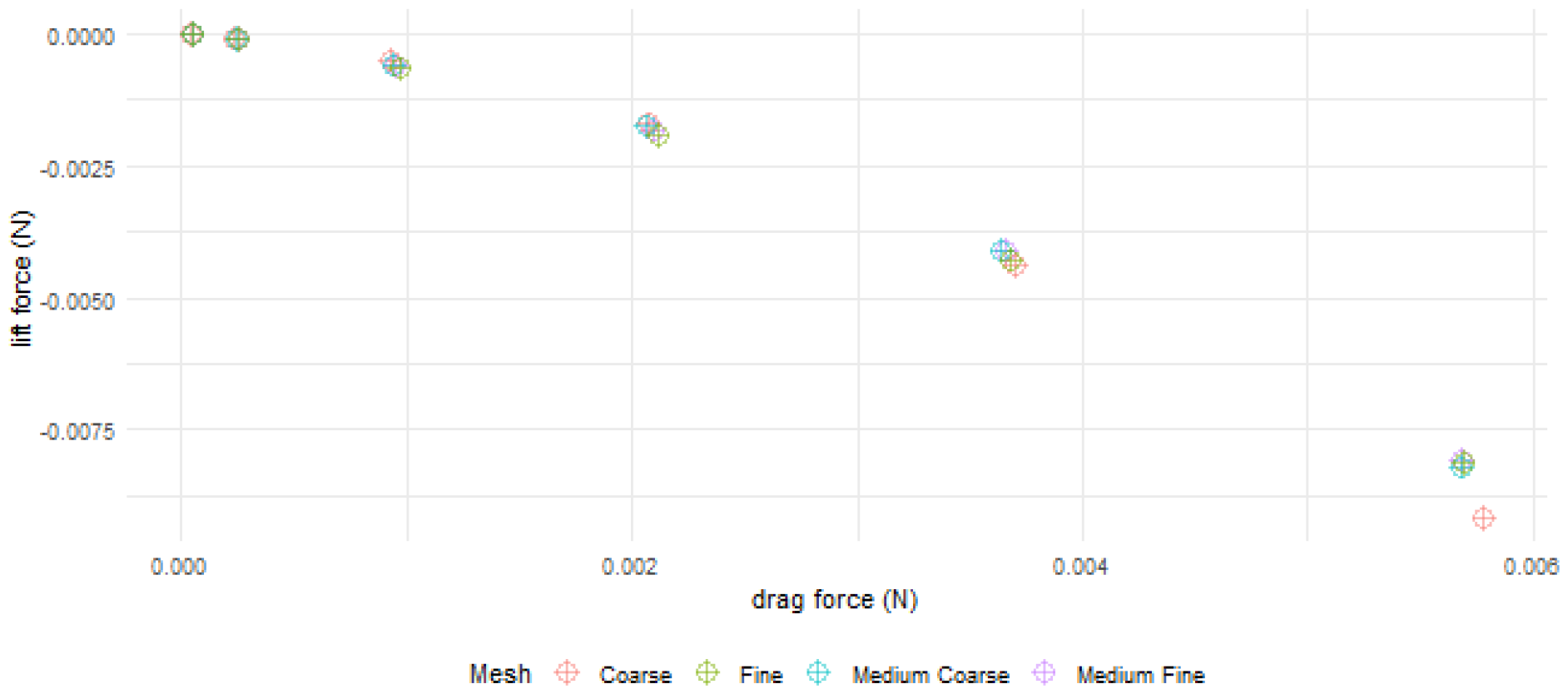
Results of CFD simulations interrogating the mesh refinement. Meshing parameters for each refinement provided in Supplementary Table 1, raw values for each simulation provided in Excel Spreadsheet uploaded to OSF (doi.org/10.17605/OSF.IO/2MAFU).

**Supplementary Table 1.** Meshing parameters used for mesh refinement analyses.

| Mesh type | Base element size (m) | Body of influence (m) | Face sizing (m) | Inflation layers # | # total elements |
| --- | --- | --- | --- | --- | --- |
| Medium-fine | $7.5 \times 10^{-3}$ | $1.7 \times 10^{-3}$ | $2.5 \times 10^{-4}$ | 20 | $2.5 \times 10^6$ |
| Coarse | $1 \times 10^{-2}$ | $3 \times 10^{-3}$ | $4 \times 10^{-4}$ | 20 | $1.6 \times 10^6$ |
| Medium-coarse | $8.5 \times 10^{-2}$ | $2 \times 10^{-3}$ | $3 \times 10^{-4}$ | 20 | $2.1 \times 10^6$ |
| Fine | $5 \times 10^{-3}$ | $1.5 \times 10^{-3}$ | $2 \times 10^{-4}$ | 20 | $3.0 \times 10^6$ |

